# Daam2 Driven Degradation of VHL Promotes Gliomagenesis

**DOI:** 10.1101/140277

**Authors:** Wenyi Zhu, Saritha Krishna, Cristina Garcia, Chia-Ching John Lin, Ken Scott, Carrie A Mohila, Chad J Creighton, Seung-Hee Yoo, Hyun Kyoung Lee, Benjamin Deneen

## Abstract

Von Hippel-Landau (VHL) protein is a potent tumor suppressor regulating numerous pathways that drive cancer, but mutations in VHL are restricted to limited subsets of malignancies. Here we identified a novel mechanism for VHL suppression in tumors that do not have inactivating mutations. Using developmental processes to uncover new pathways contributing to tumorigenesis, we found that Daam2 promotes glioma formation. Protein expression screening identified an inverse correlation between Daam2 and VHL expression across a host of cancers, including glioma. These *in silico* insights guided corroborating functional studies, which revealed that Daam2 promotes tumorigenesis by suppressing VHL expression. Furthermore, biochemical analyses demonstrate that Daam2 associates with VHL and facilitates its ubiquitination and degradation. Together, these studies are the first to define an upstream mechanism regulating VHL suppression in cancer and describe the role of Daam2 in tumorigenesis.

**Statement of Significance:** We found that the glial developmental factor Daam2 promotes glioma tumorigenesis by suppressing VHL expression. Our studies show, for the first time, a regulatory mechanism that operates upstream of VHL in cancer and provides an explanation for how VHL expression is extinguished in tumors that do not have inactivating mutations.

## Introduction

Tumor suppressor and oncogenic pathways function in part by subverting existing cellular programs to promote cancer “hallmark” properties that engender malignant growth (Hanahan and Weinberg, 2011). This corruption of normal cellular physiology is mediated by aberrant activities of these tumorigenic pathways, which are predominantly driven by genetic mutation. Importantly, genetic mutation is not the sole source of this dysregulation, as changes in gene expression via promoter methylation or protein turnover can phenotypically resemble driver mutations and contribute to tumorigenesis (Esteller et al., 1999; Hegi et al., 2004; Pineda et al., 2015; Reinstein and Ciechanover, 2006; Semenza, 2003; Shen et al., 2005; Zochbauer-Muller et al., 2001). While these broad regulatory processes have been linked to cancer, the underlying molecular mechanisms that regulate expression of key components of tumorigenic pathways are very poorly characterized.

*VHL* is a key tumor suppressor that is mutated in Von Hippel-Landau disease, a hereditary form of clear-cell renal carcinoma (ccRCC) (Chen et al., 1995; Gossage et al., 2015; Kim and Kaelin, 2004; Maher et al., 1990). VHL functions by binding to HIF1α and pAkt and modulating their degradation and activity, respectively. Mutant forms of *VHL* associated with ccRCC are incapable of binding HIF1α or pAkt, resulting in stabilized expression and activation of these proteins, which ultimately facilitates tumorigenesis (Guo et al., 2016; Ivan et al., 2001; Jaakkola et al., 2001; Maxwell et al., 1999; Min et al., 2002; Ohh et al., 2000). While dysregulated HIF1α and pAkt are associated with most forms of cancer, mutations in VHL are found predominately in ccRCC. This dichotomy suggests that additional regulatory mechanisms oversee VHL dysregulation or inactivation in other malignancies. Indeed recent studies have shown that ID2 can interfere with VHL activity in glioma cell lines (Lee et al., 2016). However, the upstream mechanisms that directly regulate *VHL* expression and protein turnover in cancer remain undefined.

One potential mode of tumor suppressor gene regulation is through developmental mechanisms. Developmental processes directly contribute to all forms of malignancy and are utilized by tumorigenic pathways to maintain cells in an undifferentiated and proliferative state (Jackson et al., 2006; Kesari et al., 2005; Stiles and Rowitch, 2008). Given these established molecular and functional interactions, it stands to reason that expression of tumorigenic pathways may be reciprocally regulated by developmental mechanisms. However, whether such reciprocal regulation of tumor suppressor pathways by developmental factors contributes to tumorigenesis is poorly defined.

To investigate the interface between developmental programs and the regulation of tumor suppressor pathways, we used malignant glioma as a model system. As a molecular entry point for these studies, we focused on Daam2, a key developmental regulator that suppresses glial differentiation and also contributes to dorsal patterning in the developing CNS (Lee et al., 2015; Lee and Deneen, 2012). Here we found that Daam2 promotes tumorigenesis in mouse and human models of malignant glioma. Bioinformatics analysis revealed that Daam2 and VHL expression is inversely correlated across a host of human malignancies. These *in silico* observations are corroborated by *in vivo* functional studies, which revealed that Daam2 promotes tumorigenesis by suppressing VHL expression. Mechanistically, we found that Daam2 associates with VHL and facilitates its degradation by the ubiquitin pathway. Together, these studies represent the initial characterization of Daam2 function in glioma and define, for the first time, an upstream regulatory mechanism that controls VHL protein expression in cancer. Moreover, because mutations in VHL are restricted to a limited set of malignancies, we have identified a new mechanism for VHL inactivation in tumors that do not have inactivating mutations.

## Results

### Daam2 is expressed in human and mouse glioma

Previously, we identified Daam2 as a component of the Wnt receptor complex that directly contributes to dorsal patterning in the embryonic spinal cord and oligodendrocyte differentiation during development and after injury (Lee et al., 2015; Lee and Deneen, 2012). To further explore its role in neurological diseases, we sought to investigate its role in malignancies of the CNS. Towards this we took advantage of the TCGA pan-cancer expression data set (Akbani et al., 2014; Cancer Genome Atlas Research et al., 2013) and evaluated Daam2 expression across a spectrum of 34 malignancies, finding that it’s most highly expressed in low-grade glioma (LGG) and glioblastoma multiforme (GBM) (Figure 1A).

**Figure 1.**
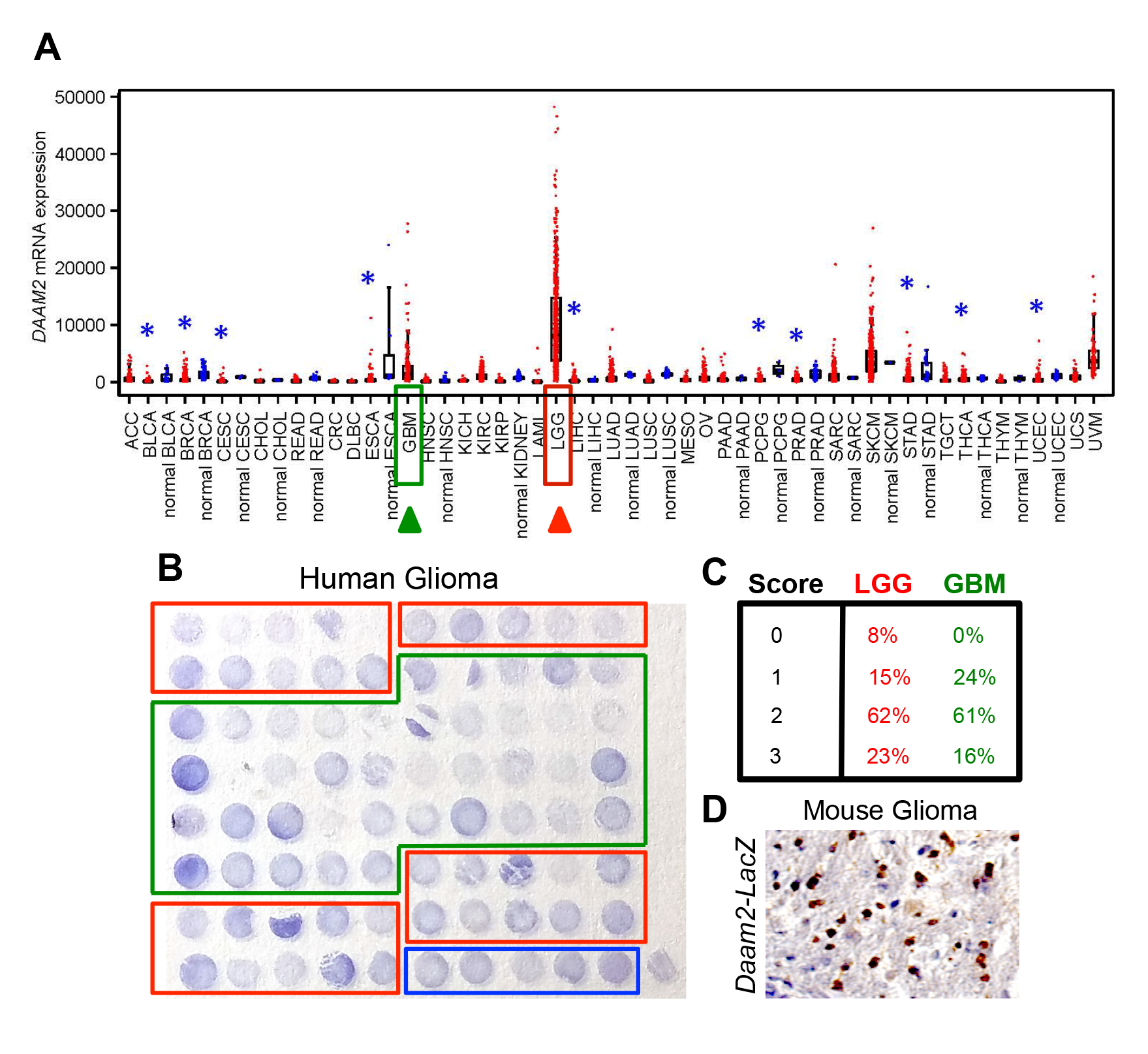
Daam2 is expressed in human glioma. (A) Analysis of Daam2 RNA expression across a spectrum of cancers. Data was generated by TCGA Research Network and is publicly available (see methods). Box plots represent 5%, 25%, 50%, 75%, and 95%. Asterisk denotes tumor groups significantly higher (red) or lower (blue) as compared to corresponding non-tumor (“normal”) group. P-values were generated by t-test on log-transformed data. (B) Tissue microarray showing in situ hybridization expression analysis of human Daam2 across 35 LGG (red boxes) and 40 GBM (green boxes). Blue box denotes normal human brain samples. (C) Graded pathological scoring (0-low intensity, 3-high intensity) of Daam2 expression in the samples in B; values are the percentage of LGG and GBM tumors from the tissue microarray that were assigned respective score. (D) Immunohistochemistry analysis of Daam2-LacZ expression in mouse glioma generated in Daam2^LacZ/+^ mice.

To validate Daam2 expression in LGG and GBM, we used in situ hybridization (ISH) across a cohort of 35 LGG and 40 GBM primary human samples, finding that Daam2 demonstrates heterogeneous expression within each glioma sub-type (Fig.1B-C). Notably, the majority of tumors exhibited staining intensity scores that exceeded normal brain samples, indicating that Daam2 expression is elevated within glioma tumors (Fig. 1B-C; Supplemental Figure S1). Next, we evaluated Daam2 expression in our mouse model of malignant glioma, where we combine in utero electroporation (IUE), with CRISPR-mediated deletion of *NFI, PTEN,* and *p53* (herein CRISPR/IUE) (Supplemental figure S2) (Chen et al., 2016; Chen and LoTurco, 2012; Chen et al., 2012; Cong et al., 2013; John Lin et al., 2017b). This model closely resembles the genetics of GBM (Alcantara Llaguno et al., 2009; Kwon et al., 2008; Zhu et al., 2005) and begins producing tumors detectable around post-natal week 8. Combining this model, with our the Daam2-LacZ mouse line, we performed immunostaining on the resultant tumors, finding that Daam2 exhibits elevated expression levels in tumors, compared to normal brain tissue (Figure 1D). Finally, we evaluated Daam2 expression in xenograft tumors generated from primary human GBM cell lines, finding that Daam2 is also highly expressed in these human cell line models (Supplemental Figure S1). Put together, these studies indicate that Daam2 expression is elevated in both human LGG and GBM and is expressed in the associated model systems.

### Overexpression of Daam2 accelerates glioma tumorigenesis

The forging expression analysis in human glioma and associated mouse models led us to examine whether Daam2 contributes to glioma tumorigenesis. To assess its role in glioma, we performed overexpression, gain-of-function (GOF) studies in human GBM cell lines, finding that Daam2 accelerates the rate of cell growth *in vitro* (Figure 2A-B). Next, we determined how Daam2 influences anchorage independent growth, via soft agar assay, finding that it also accelerates colony formation (Figure 2C-D). Together, these *in vitro* studies, indicate that Daam2 promotes cell proliferation and growth in human GBM cell lines

**Figure 2.**
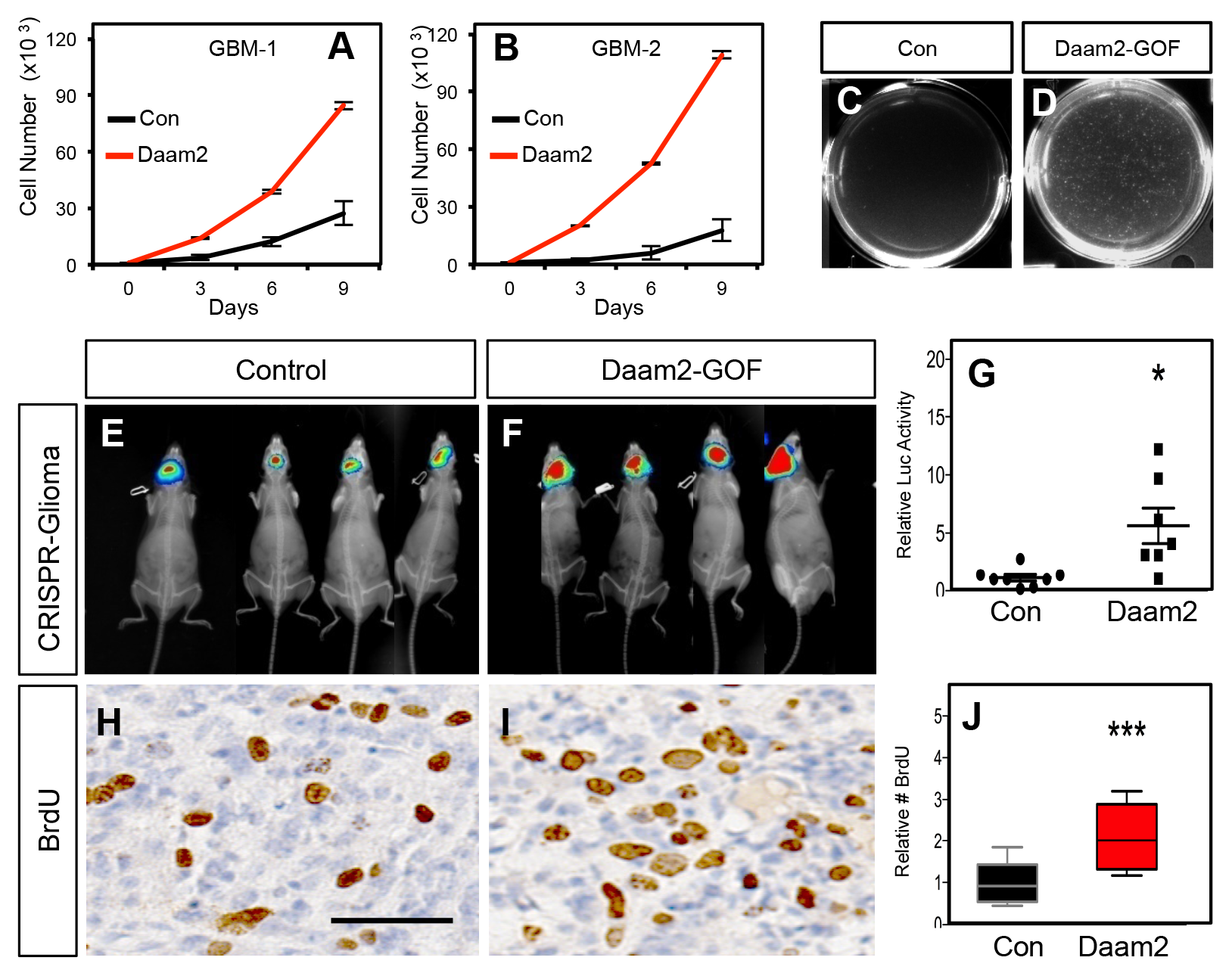
Daam2 accelerates glioma tumorigenesis. (A-B) Cell proliferation analysis of human GBM1 and GBM2 cell lines infected with lentivirus containing Daam2 or GFP control. (C-D) Soft agar assay with GBM1 cell line infected with lentivirus containing Daam2 or GFP control, images are representative. Each of these *in vitro* experiments was performed in triplicate and replicated three times. (E-G) Representative bioluminescence imaging of mice bearing CRISPR-IUE glioma with Daam2 overexpression or control, imaged at 7 weeks of age. Imaging quantification in G is derived from 7 Daam2-overexpression and 8 control mice; *p=0.0079. (H-J) Immunohistochemistry analysis of BrdU expression from mice bearing CRISPR-IUE glioma from each experimental condition. Relative number of BrdU expressing cells is quantified in J and is derived from 6 total mice, 4 slides per mouse, from each experimental condition; ***p=0.0019. Scale bar in H is 50nm. Error bars in G and J are +/− SEM.

To evaluate the role of Daam2 in tumorigenesis, we turned to mouse models of malignant glioma. The first model combines targeted PiggyBac overexpression of oncogenic Ras-V12 (PB-Ras) in glial precursors with IUE, to generate malignant glioma in mice around post-natal day 14 (Supplemental Figure S2). Combining PiggyBac-mediated overexpression of Daam2 with this Ras-driven model resulted in an acceleration of tumorigenesis (Supplemental Figure S3). To further substantiate these findings in additional mouse models, we next used our CRISPR/IUE model (Supplemental Figure S2). Consistent with the Ras model, combined overexpression of Daam2 in the CRISPR/IUE model, also resulted in accelerated tumorigenesis (Figure 2E-G). BrdU labeling analysis of the resulting tumors from both models revealed that overexpression of Daam2 results in an increase in the number BrdU-expressing cells (Figure 2H-J; Supplemental Figure S3). These data, combined with our *in vitro* studies, indicate that overexpression of Daam2 promotes glioma cell proliferation and tumorigenesis.

### Loss of Daam2 impairs glioma tumorigenesis

To further delineate the role of Daam2 in glioma, we next performed a series of complementary loss-of-function (LOF) studies in human and mouse glioma models. In human GBM cell lines we performed shRNAi-mediated knockdown of human Daam2, finding that decreased expression of Daam2 inhibited their rate of growth *in vitro* (Figure 3A-B). Next, we assessed the tumorigenic potential of these GBM cell lines, finding that knockdown of Daam2 resulted in a significant decrease in tumor growth *in vivo* (Figure 3C-E). These knockdown studies across both *in vitro* and *in vivo* systems, complement the overexpression studies, and further substantiate the role of Daam2 in glioma cell proliferation and tumorigenesis.

**Figure 3.**
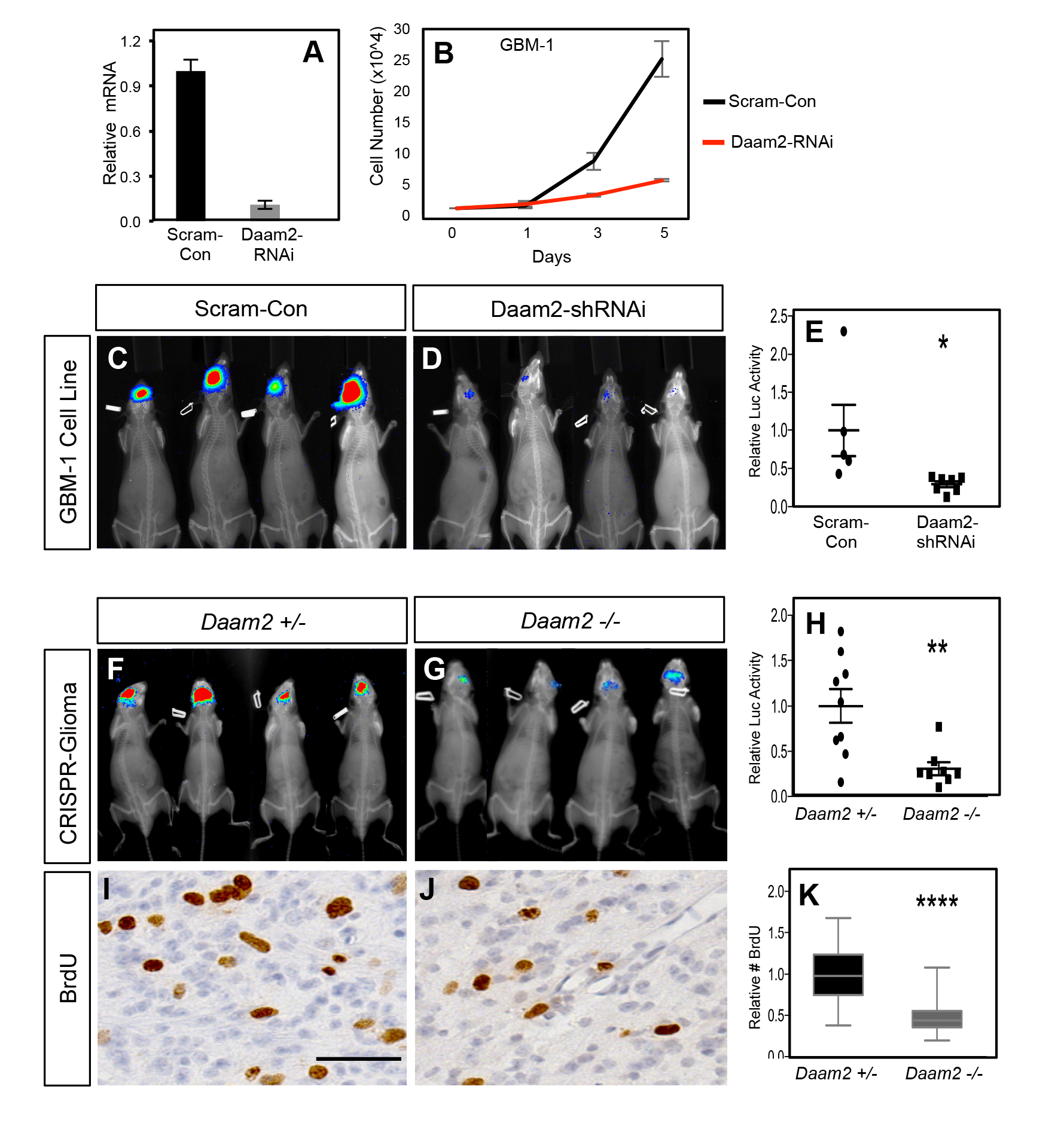
Loss of Daam2 suppresses glioma tumorigenesis. (A) qRT-PCR demonstrating effective knockdown of human Daam2 mRNA expression in human GBM1 cell line infected with lentivirus containing Daam2-shRNAi or scrambled control. (B) Cell proliferation analysis of human GBM1 cell line infected with human Daam2-shRNAi or scrambled control lentivirus. (C-E) Representative bioluminescence imaging of mice transplanted with GBM1 cell lines transfected with Daam2-shRNAi or scrambled control, imaged 6 weeks post-transplantation. Imaging quantification is derived from 7 mice transplanted with GBM1/Daam2-shRNAi and 5 GBM1/scrambled controls; *p=0.0325. (F-H) Representative bioluminescence imaging of mice bearing CRISPR-IUE glioma generated in *Daam2*+/− or *Daam2*−/− mice, imaged at 8 weeks of age. Imaging quantification is derived from 8 *Daam2*−/− and 9 *Daam2*+/− mice; **p=0.0049. (I-K) Immunohistochemistry analysis of BrdU expression from mice bearing CRISPR-IUE glioma from each experimental condition. Relative number of BrdU expressing cells is quantified in K and is derived from 6 total mice, 4 slides per mouse, from each experimental condition; ****p=0.00001. Scale bar in I is 50nm. Error bars in E, H and K are +/− SEM.

To further evaluate the necessity for Daam2 in glioma tumorigenesis, we used CRISPR/IUE model to generate malignant glioma in *Daam2*^+/−^ and *Daam2*^−/−^ mice (Lee et al., 2015). As shown in Figure 3F-H, mice lacking Daam2 demonstrated decreased rates of tumor formation in this model compared to the heterozygote control. Moreover, BrdU labeling revealed substantial decreases in the number of proliferating cells in *Daam2*^−/−^ tumors (Figure 3I-K). Together, our LOF, and complementing GOF (Figure 2) studies in human and mouse models of glioma indicate that Daam2 promotes cell proliferation and tumorigenesis.

### Daam2 suppresses VHL expression

Having established that Daam2 functions to promote glioma tumorigenesis, we next sought to uncover the mechanism by which it operates. Previously, we found that Daam2 functions as a positive regulator of Wnt-signaling in the developing CNS (Lee and Deneen, 2012), suggesting that it may also function in this manner in glioma. To evaluate Wnt-activity we used an established Wnt-reporter (TOP-FLASH) and found that modulation of Daam2 expression has a modest effect on Wnt activity in glioma cell lines and did not impact Wnt activity in our mouse model of glioma (Supplemental Figure S4).

That Wnt-activity is not affected by changes in Daam2 expression raises the question of how Daam2 promotes glioma tumorigenesis. To identify changes in protein expression associated with Daam2-mediated tumorigenesis we performed reverse phase protein lysate microarray (RPPA) on FACS-isolated mouse glioma samples that overexpress Daam2. Analysis revealed a cohort of proteins that are strongly downregulated in glioma samples that overexpress Daam2 (Figure 4A; Supplemental Figure S4; Supplemental Table I). To further substantiate these potential relationships, we leveraged existing TCGA RNA-Seq and RPPA data to determine whether there is an inverse correlation between Daam2 expression and this cohort of proteins across a spectrum of human malignancies (Supplemental Figure S5). This analysis identified Von Hippel Landau (VHL) as one of the proteins from this cohort with the most significant inverse correlation score across this spectrum of malignancies, with lung adenocarcinoma and GBM demonstrating the strongest negative correlations between Daam2 and VHL (Figure 4B; Supplemental Figure S5).

**Figure 4.**
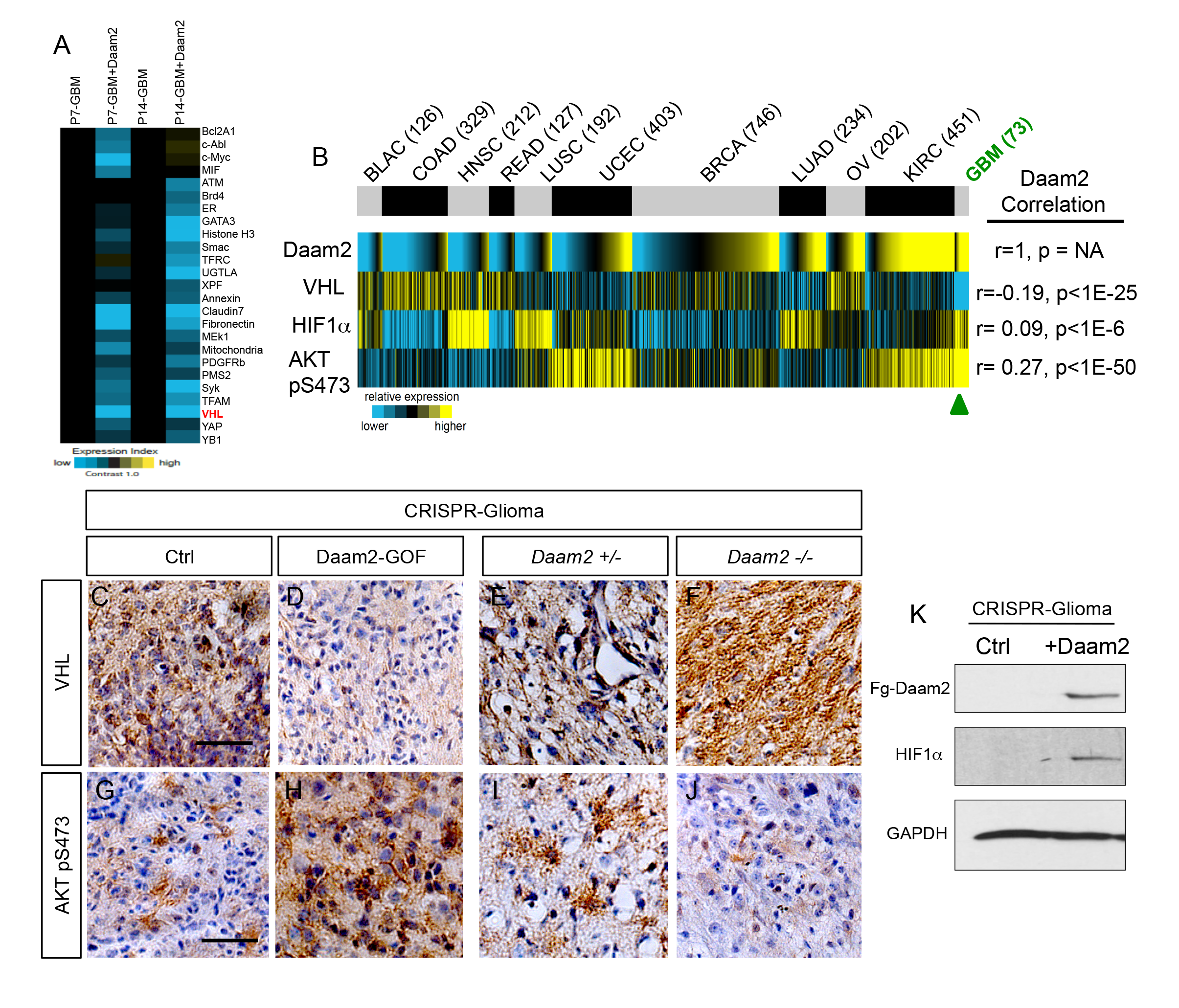
Daam2 suppresses VHL expression. (A) Heatmap analysis of RPPA data showing the core cohort of proteins downregulated by overexpression of Daam2 in the PB-Ras mouse glioma model. (B) Correlation of Daam2 mRNA expression with VHL and Akt pS473 protein expression data and HIF1α mRNA expression. Protein and mRNA expression data from samples was obtained from The Cancer Proteome Atlas (TCPA) and correlations with Daam2 mRNA was performed using Pearson’s. Abbreviated cancer type is listed on top panel and number of tumors analyzed in the TCPA dataset for each cancer type is listed in parenthesis; GBM is denoted by green text. (C-F) Representative immunohistochemical analysis of VHL expression in CRISPR-IUE glioma tumors derived from Daam2-overexpression (D) or knockout of Daam2 (F) and associated controls. (G-J) Representative immunohistochemical analysis of Akt pS473 expression in CRISPR-IUE glioma tumors derived from Daam2-overexpression (H) or knockout of Daam2 (J) and associated controls. Images are representative of analysis performed on 6 independent tumors for each experimental condition. (K) Immunoblot with antibodies against HIF1α from protein lysates derived from CRISPR-IUE glioma overexpressing Daam2 or control. Scale bars in C and G are 50nm.

VHL is an established tumor suppressor that is frequently mutated in clear cell renal carcinoma (ccRCC) and functions by facilitating the degradation of HIF1α and pAkt (Guo et al., 2016; Ivan et al., 2001; Jaakkola et al., 2001; Maxwell et al., 1999; Min et al., 2002; Ohh et al., 2000). Given that Daam2 expression is inversely correlated with VHL, we next assessed whether Daam2 demonstrates congruent correlation with HIF1α and pAkt in the panel of human cancers. Analysis of these data revealed that Daam2 expression was positively correlated with HIF1α and pAkt protein expression across this cohort of malignancies, with GBM and lung adenocarcinoma also showing the strongest positive correlation (Figure 4B). Together, our RPPA screen and associated bioinformatics analysis of human malignancies indicate that Daam2 expression is inversely correlated with VHL expression and positively correlated with its downstream signaling axis.

### Daam2 promotes tumorigenesis through VHL

The RPPA data suggests that Daam2 modulates expression of VHL in glioma. To directly test this hypothesis, we evaluated expression of VHL and its signaling axis in our Daam2-GOF and Daam2-LOF mouse tumor models. Immunostaining for VHL in these models corroborated our RPPA data, where VHL expression was dramatically reduced in Daam2-GOF tumors, and increased in the *Daam2*^−/−^ tumors (Figure 4C-F). Next we assessed expression of VHL’s downstream effectors, pAkt and HIF1α, finding that both of these proteins demonstrated elevated levels of expression in the Daam2-GOF tumors, and pAkt having decreased levels in the *Daam2*^−/−^ tumors (Figure 4G-K). One consequence of elevated HIF1α is increased angiogenesis (Fan et al., 2014; Keith and Simon, 2007; Semenza, 2004), which we detected in Daam2-GOF tumors via staining for the endothelial marker, CD31 (Supplemental Figure S6). These data, combined with our bioinformatics analysis, indicate that Daam2 suppresses VHL-signaling in malignant glioma.

These observations raise the question of whether the effects of Daam2 on glioma tumorigenesis are mediated through its suppression of VHL expression. To determine whether an epistatic relationship exists, we overexpressed VHL in the presence of Daam2 overexpression in human GBM cell lines, finding that VHL suppresses the increased rate of cell growth mediated by Daam2-alone (Figure 5A-B). Next, we extended these studies to our mouse models of glioma, finding that combined overexpression of VHL with Daam2 similarly suppressed the accelerated rate of tumorigenesis and proliferation mediated by Daam2-alone (Figure 5C-I). Moreover, overexpression of VHL resulted in concordant suppression of pAkt and angiogenesis in these tumors (Figure 5J-P; Supplemental Figure S6). Put together, these data indicate that Daam2 promotes tumorigenesis by suppressing VHL expression in glioma.

**Figure 5.**
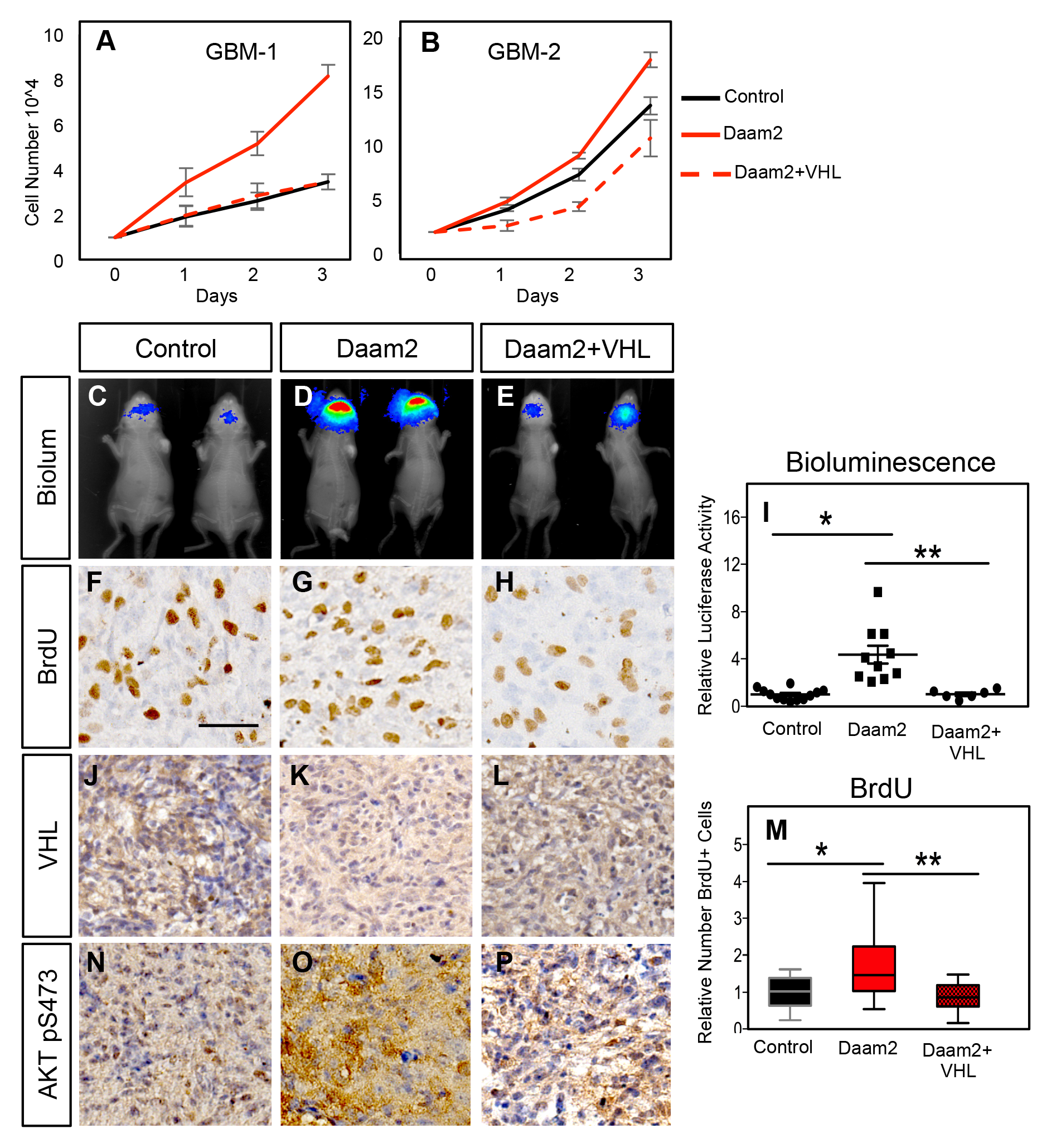
Daam2 promotes gliogenesis via suppression of VHL. (A-B) Cell proliferation analysis of human GBM1 and GBM2 cell lines infected with lentivirus containing Daam2, GFP control, or Daam2+VHL. (C-E, I) Representative bioluminescence imaging of mice bearing PB-Ras glioma with Daam2 overexpression, Daam2+VHL overexpression, or control, imaged at 10days post-natal. Imaging quantification in I is derived from 10 Daam2-overexpression, 6 Daam2+VHL, and 12 control mice; *p=0.0001, **p=0.0046. (F-H, M) Immunohistochemistry analysis of BrdU expression from mice bearing PB-Ras glioma from each experimental condition. Relative number of BrdU expressing cells is quantified in M and is derived from 6 total mice, 4 slides per mouse, from each experimental condition; *p=0.0089, **p=0.0007. (J-L) Representative immunohistochemical analysis of VHL expression in PB-Ras glioma from described experimental conditions and associated controls. (N-P) Representative immunohistochemical analysis of Akt pS473 expression in PB-Ras glioma tumors derived from the described experimental conditions and associated controls. Images in J-P are representative of analysis performed on 6 independent tumors for each experimental condition. Statistics derived by one-way analysis of variance (ANOVA) and followed by Tukey’s test for between-group comparisons. Scale bar in F is 50nm. Error bars in I and M are +/− SEM.

### Daam2 facilitates ubiquitination of VHL protein

Analysis of human cancer data, along with our functional studies in mouse models, strongly suggests that Daam2 expression results in the loss of VHL protein. To determine the mechanism by which Daam2 influences the levels of VHL protein, we next performed a series of immunoprecipitation experiments to evaluate the biochemical relationship between Daam2 and VHL. Using protein lysates extracted from our PB-Ras model of glioma, we found that Daam2 co-immunoprecipitates with VHL, suggesting that these proteins associate (Figure 6A).

**Figure 6.**
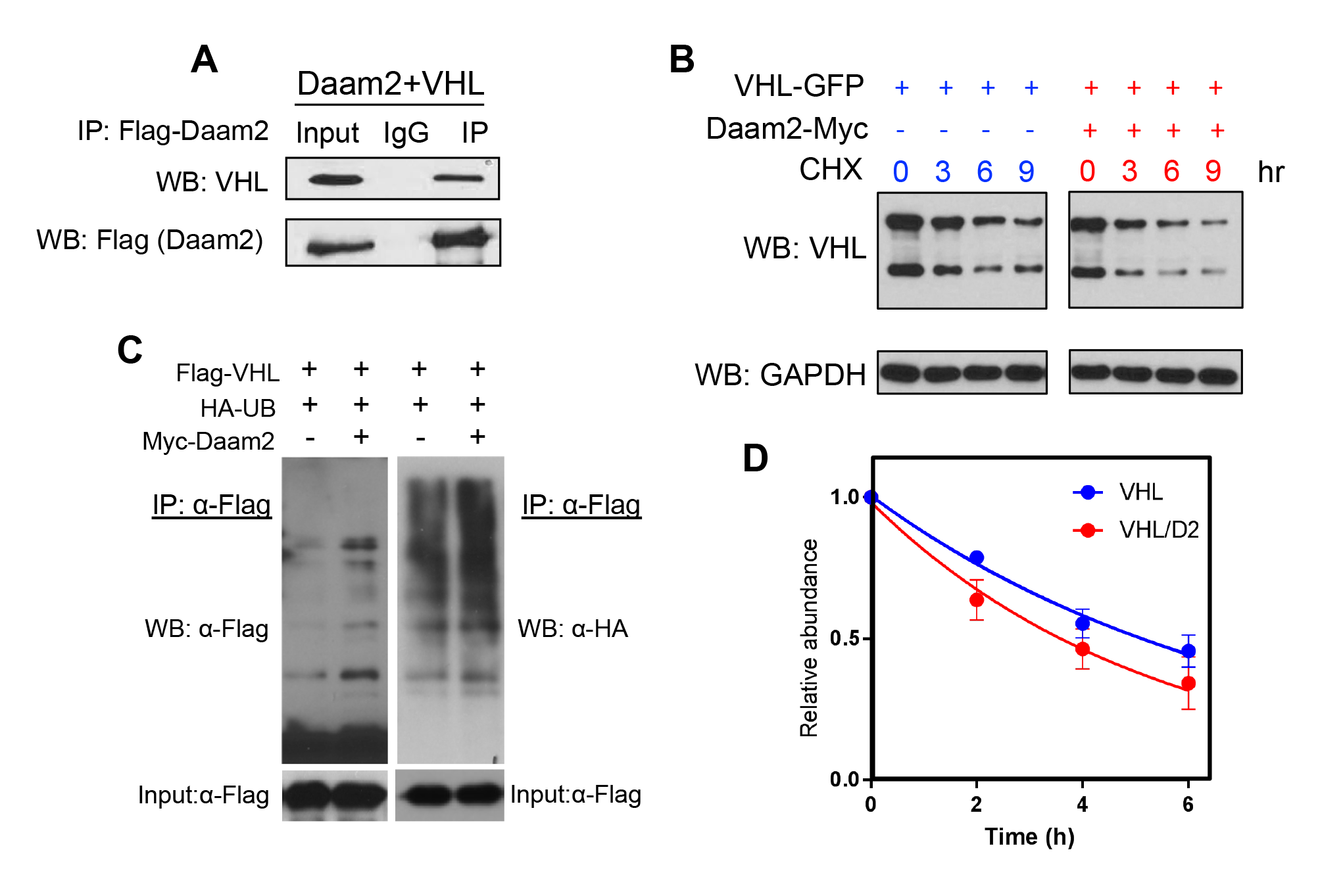
Daam2 mediates VHL degradation and ubiquitination. (A) Immunoprecipitation of Flag-Daam2, followed by immunoblot with VHL on lysates derived from PB-Ras glioma overexpressing Flag-Daam2. Detection of endogenous VHL in the “IP” lane, indicates Daam2 associates with VHL in mouse glioma. (B,D) 293 cells were transfected with VHL and Daam2 (or control). 24 hrs after transfection, cells were treated with 40μg/ml cycloheximide and incubated for the indicated time before harvest. Western blotting was performed to monitor VHL levels using anti-VHL antibody. Quantification in D shows the effect of Daam2 on VHL stability. Half-life of VHL (5.082hr) was significantly decreased in the presence of Daam2 (3.679hr). Error bars represent mean ± SEM (n=3). Half-life was determined via nonlinear, one-phase exponential decay analysis (half-life parameter, K, is significantly different in two conditions: p=0.0081). Error bars represent mean ± SEM (n=3). (C) Daam2 enhances VHL ubiquitination. 293T cells were co-transfected with Flag-VHL, HA-UB(ubiquitin), and Myc-Daam2. Cells were treated with MG132 (10ug/ml) 6 hrs before harvest. Whole-cell lysate were immunoprecipitated with anti-Flag M2 beads then analyzed by western blotting with Flag and HA antibodies.

Given that Daam2 associates with VHL, and expression of VHL protein is negatively correlated with Daam2, we hypothesized that these expression dynamics are the result of Daam2 promoting the degradation of VHL. To test this we co-transfected Daam2 and VHL in 293 cells, and measured the rate of cyclohexamide-mediated degradation, finding that VHL degradation is significantly enhanced in the presence of Daam2 (Figure 6B, D). The ubiquitination pathway is a central mechanism that oversees protein degradation, (Pickart and Eddins, 2004) (Kerscher et al., 2006; Ulrich and Walden, 2010), therefore, we next examined whether Daam2 facilitates the ubiquitintion of VHL. Using *in vitro* systems, we found that in the presence of elevated levels of Daam2, the extent of VHL ubiquitination is substantially increased and levels of VHL protein demonstrate a concomitantly decrease (Figure 6C). Together, these mechanistic studies reveal that the underlying biochemical relationship between Daam2 and VHL is mediated by ubiquitin-driven protein degradation.

## Discussion

Using factors critical for central nervous system (CNS) development as an entry point to identify new mechanisms that contribute to tumorigenesis, we found that the glial development factor, Daam2 promotes glioma tumorigenesis across human glioma cell lines and multiple mouse models of glioma. Protein screening and bioinformatics studies revealed that Daam2 and VHL expression is inversely correlated across a broad spectrum of cancers, while functional interrogation of this relationship demonstrated that Daam2 promotes tumorigenesis via suppression of VHL expression. Mechanistically, Daam2 associates with VHL and stimulates its ubiquitination and degradation. Together, these studies are the first to define an upstream mechanism regulating VHL protein turnover in cancer and describe the role of Daam2 in tumorigenesis.

### Daam2 stimulates glioma tumorigenesis

Leveraging existing TCGA expression data across a broad spectrum of malignancies, we found that Daam2 is most highly expressed in glioma and melanoma. These expression characteristics in glioma were confirmed using tissue arrays and functional studies revealed that Daam2 promotes cell proliferation and tumorigenesis in human and mouse glioma models. These are the first studies to describe the role of Daam-family proteins in tumorigenesis. Daam1 and Daam2 have highly conserved formin domains yet exhibit nono-verlapping expression patterns in the developing CNS (Habas et al., 2001; Kida et al., 2004; Lee and Deneen, 2012; Nakaya et al., 2004), suggesting distinct functions during development. Indeed, Daam2 functions in this context via canonical Wnt-signaling, while Daam1 operates via the non-canonical Wnt, planar cell polarity (PCP) pathway (Habas et al., 2001; Lee and Deneen, 2012; Li et al., 2011; Liu et al., 2008; Zhu et al., 2012). It will be important to discern whether this functional diversity between Daam1 and Daam2 is also applicable to CNS malignancies. Given that the PCP pathway has been implicated in tumor cell invasion and migration (Anastas and Moon, 2013; Paw et al., 2015; Weeraratna et al., 2002), it is possible that Daam-family proteins also contribute to these features of tumorigenesis. Recent studies have implicated Daam1, Daam2 and other formin-family genes in the migration of breast and neuroblastoma cell lines (Luga et al., 2012), respectively, suggesting that the Daam-family proteins may also contribute to invasion and metastasis in other malignancies.

Our previous studies have shown that Daam2 potentiates canonical Wnt signaling through the clustering of existing Wnt receptor complexes (Lee and Deneen, 2012). While the Wnt pathway has been implicated in several forms of cancer, including medulloblastoma, mutations in key components of the Wnt-pathway and dysregulated Wnt-signaling has not been widely linked to low- or high-grade glioma (Bienz and Clevers, 2000; de La Coste et al., 1998; Klaus and Birchmeier, 2008; Lustig and Behrens, 2003; Morin et al., 1997; Rubinfeld et al., 1997; Zurawel et al., 1998). This, coupled with the fact that Daam2 acts upon existing Wnt receptor complexes, may explain why we did not witness any overt changes in Wnt activity when we manipulated Daam2 expression in our glioma models. Nevertheless, we cannot formally rule out a possible role for Daam2 in canonical or non-canonical Wnt signaling in glioma or in other malignancies driven by Wnt dysregulation.

### New Parallels Between Development and Cancer

Our observation that manipulation of Daam2 expression promoted tumorigenesis, but not canonical Wnt signaling, points to the possibility that it functions through a Wnt-independent mechanism. RPPA and bioinformatics approaches revealed that Daam2 expression is inversely correlated with a cohort of genes (Figure S5), including VHL, across a broad spectrum of cancers. Moreover, the RPPA data analysis identified a larger cohort of proteins that were downregulated in the presence of Daam2 in our mouse model of glioma. These observations, coupled with our findings that Daam2 associates with- and facilitates- the ubiquitination of VHL, suggest that it may also play a role in the ubiquitination pathway. The nature of this prospective relationship between Daam2 and the existing ubiquitination machinery, and whether these relationships are specific to malignancy or can be extended to development are areas of future investigation and represent potentially new parallels between development and cancer. Indeed dysregulation of ubiquitination is linked to protein aggregation associated with several neurological disorders, functioning through both neuronal and glial populations (Jansen, et al. 2014 and Hedge and Upadhya, 2007)

Another feature of Daam2 is that it functions to suppress the differentiation of oligodendrocyte progenitor cells (OPC) during development and after injury (Lee et al., 2015). Studies in mouse models suggest that OPCs may serve as a cell of origin for glioma, while NG2-positive OPC populations are endowed with tumor initiating properties (Ligon et al., 2007; Liu et al., 2011; Persson et al., 2010; Stiles and Rowitch, 2008; Sugiarto et al., 2011; Yadavilli et al., 2015). Moreover, OPCs have latent proliferative capacity that is essential for injury responses, suggesting parallels between the processes that drive injury responses and tumorigenesis. Indeed recent studies have linked the VHL-HIF1α signaling axis to OPC development and neonatal hypoxic injury responses (Yuen et al., 2014). That Daam2 promotes tumorigenesis via suppression of VHL and also regulates OPC development and injury responses, suggests that it may also utilize these tumorigenic mechanisms in the context of injury. Together these findings add to the emerging evidence that OPC associated signaling networks play critical roles in glioma pathophysiology and further reinforce the parallels between neurological disorders and cancer biology.

### Regulation of VHL in cancer

Mechanistic studies revealed that Daam2 promotes glioma tumorigenesis via suppression of VHL, a classic tumor suppressor that promotes HIF1α degradation and inhibits Akt activity. Importantly, inactivating mutations in VHL are predominately found in ccRCC and are very rare in most other forms of cancer, including glioma. These observations raise the critical question of how HIF1α becomes upregulated in cancers that do not exhibit VHL mutations. Our studies show, for the first time, a regulatory mechanism that operates upstream of VHL in cancer and provides an explanation for how VHL expression is extinguished in tumors that do not have inactivating mutations. Moreover, because Daam2-VHL expression demonstrates a robust, inverse correlation across a spectrum of cancers, this is likely to be a generalized mechanism of VHL suppression in cancer. Interestingly, prior studies in yeast have shown that VHL can be degraded via the Hsp70 and Hsp110 chaperone system (McClellan et al., 2005; Melville et al., 2003). Given that Hsp’s are expressed across a host of cancers (Garcia-Morales et al., 2007; Nylandsted et al., 2002; Sauvageot et al., 2009), it will be important to determine whether Daam2 engages the Hsp chaperone system to facilitate VHL degradation in glioma and other malignancies.

Given its role as the central regulator of HIF1α, surprisingly little is known about the regulatory biology surrounding VHL and its role in glioma formation. Studies in glioma cell lines suggest that ID2 interference with VHL activity can deregulate HIF1α expression and promote tumorigenesis in xenograft models (Lee et al., 2016). Interestingly, ID2 has also been linked to the suppression of OPC differentiation via direct regulation of cell cycle kinetics (Wang et al., 2001). These observations further reinforce the relationship between OPC development and glioma tumorigenesis and highlight key parallels between ID2 and Daam2 function across these systems, as both genes suppress VHL, inhibit OPC differentiation, and promote glioma tumorigenesis. Because both proteins associated with- and regulate-VHL, albeit via distinct mechanisms, it’s possible that these suppressive functions are coordinated and contribute tumorigenesis. From this, a model emerges where ID2 displaces VHL from the ubiquitin complex, which facilitates its association with Daam2 and its subsequent ubiquitination and degradation. It will be important to decipher whether ID2 and Daam2 functions are in fact coordinated, and the extent to which Hsp proteins participate in this mechanism. Finally, understanding whether and how these prospective relationships are also applicable to OPC development and injury responses may also reveal new insights in CNS repair mechanisms.

## Materials and Methods

### TCGA database analysis and Comparative Bioinformatics

RNA expression data underlying the results presented in Figure 1 were generated by TCGA Research Network (http://cancergenome.nih.gov/). All data used in this study were publicly available, e.g. from The Broad Institute’s Firehose pipeline (http://gdac.broadinstitute.org/). From TCGA, we collected molecular data on 10224 tumors of various histological subtypes, for which RNA-seq data (v2 platform alignment) were available. Correlation between VHL and AKT pS473 protein on TCGA samples, were obtained from The Cancer Proteome Atlas, or TCPA.

### In Utero Electroporation (IUE)

To generate mouse gliomas, we performed in utero electroporation (IUE). Briefly, the uterine horns were exposed, and DNA combination was injected into the embryonic lateral ventricles along with Fast Green dye as the indicator. Then electroporation was accomplished by BTW tweezertrodes connected with the pulse generator (BTX 8300) in the setting of 33V, 55ms per pulse100ms intervals. In CRISPR-IUE model, the DNA combination is composed of the “helper-plasmid” pGLAST-PBase (2.0 ug/ul) and all the other DNA (1.0 ug/ul), including pbCAG-GFP, pbCAG-Luciferase, crNF1, crPTEN, crp53(Chen and LoTurco, 2012; John Lin et al., 2017a). In the HRas-IUE model, the DNA combination is composed of pGLAST-PBase (2.0 ug/ul) and others (1.0 ug/ul), including pbCAG-GFP2aHRas and pbCAG-Luciferase.

### Glioma Xenograft Assays

6-week-old SCID male mice (Taconic) were used for human GBM cell line transplantation. 4x10^4^ luciferase-infected primary GBM cells were injected into each mouse brain, at the location of the 1mm front, 2mm right, 3mm deep from the Bregma. Animals were euthanized and perfused six weeks after transplantation of GBM cells. Brains were fixed in 4% paraformaldehyde and 70% EtOH overnight, respectively. After fixation, brains were embedded in paraffin, sectioned and subjected to molecular and pathological analysis via immunostaining or hematoxylin and eosin staining.

### Bioluminescent Imaging

To measure the tumor growth after manipulation of Daam2 we performed bioluminescent imaging before harvesting. Mice were monitored once a week. D-Luciferin (Perkin Elmer, #122799) was diluted to 15 mg/ml with PBS and injected into each mouse at a dose of 10 ul/g body weight.

### Cell Growth Curve and Agar Assays

For *in vitro* human GBM cell line studies, adult GBM-1 and GBM-2 cell lines were kindly provided from Dr. Nabil Ahmed (Hegde et al., 2013). To manipulate the expression of Daam2 in GBM cell lines, we used the lentivirus as described above. To virally infect human primary GBM cells, 1.5x10^4^ cells were plated in 6-well plates with DMEM + 10% FBS. Cells were infected with either Daam2 or Daam2 shRNAi virus for 14 hours to make the stable cell line for GOF and LOF study, respectively. To assess rates of cell growth, 2x10^4^ cells were plated in 12-well plates and cells were counted over the course of three days.

For the agar assay, 2.5x10^4^ cells (cell agar layer) was mixed with 5ml Iscove’s 1.4% Nobel Agar + 40% FBS, covered by top and bottom coating agar layers (2ml Iscove’s 1.4% Nobel Agar + 20% FBS) in a 6mm plate. We tightly monitored the colony formation for several weeks.

### In Vitro Degradation Assay

To test VHL degradation, 1 × 10^5^ 293T cells were plated into the 12 well plate one day before transfection. 100 ng Myc-tagged Daam2 and GFP-tagged VHL was transfected by iMFectin DNA Transfection Reagent (GenDEPOT) according to the protocol. 30 hours post-transfection, cells were treated with 50 ul/ml CHX for 0, 2, 4, 6, hours. Lysis and Western blot process is described in the supplemental methods. To quantify the protein abundance the Western blot band intensity in the 0-hour sample of both VHL+/− Damm2 groups are set as “1”. The intensity of 2, 4, 6-hour degradation western blot band is normalized and plotted using Nonlinear Regression Curve by GraphPad.

### Statistical Analysis

One-way ANOVA was used to analyze bioluminescent intensity and BrdU-positive cell counts to determine the differences between Ctrl, D2 and D2/VHL groups, followed by Tukey’s test to compare between individual groups, which is demarcated by an asterisk in the graphs. Independent *t*-test was used to analyze the differences in bioluminescent intensity and BrdU-positive cell counts between Ctrl vs. D2, Het vs. KO, Scrambled-shRNAi vs. D2-shRNAi.

## Author Contributions

Conceptualization: W.Z, H.K.L, and B.D.; Methodology: W.Z., S.K., C.G., C.C.L., C.A.M., S.H.Y., H.K.L; Data Analysis: W.Z., C.J.C, C.A.M., H.K.L., B.D.; Key Reagents: K.S., S.H.Y; Writing Manuscript: W.Z., H.K.L, and B.D.

## Acknowledgements

This work was supported by grants from the Sontag Foundation (BD), Cancer Prevention Research Institute of Texas (RP510334 and RP160192 to BD and CC), and the National Institutes of Health (NS071153 and NS089366 to BD). National Multiple Sclerosis Scoiety (TA3054-A to HKL). We acknowledge the assistance of the Baylor College of Medicine Mouse Phenotyping Core, the Lester and Sue Smith Breast Center’s Pathology Core, and Functional Proteomics RPPA Core Facility at MD Anderson Cancer Center, this facility is funded by NCI # CA16672. This project was also supported in part by the IDDRC grant number 1U54 HD083092 from the Eunice Kennedy Shriver National Institute of Child Health & Human Development.

